# Assocplots: a python package for static and interactive visualization of multiple-group GWAS results

**DOI:** 10.1101/062737

**Authors:** Ekaterina A Khramtsova, Barbara E. Stranger

**Affiliations:** Department of Medicine, Section of Genetic Medicine, The University of Chicago, Chicago, IL; Institute for Genomics and Systems Biology; 3Center for Data Intensive Science, The University of Chicago, Chicago, IL

## Abstract

**Summary:** Over the last decade, genome-wide association studies (GWAS) have generated vast amounts of analysis results, requiring development of novel tools for data visualization. Quantile-quantile plots and Manhattan plots are classical tools which have been utilized to visually summarize GWAS results and identify genetic variants significantly associated with traits of interest. However, static visualizations are limiting in the information that can be shown. Here we present **Assocplots**, a python package for viewing and exploring GWAS results not only using classic static Manhattan and quantile-quantile plots, but also through a dynamic extension which allows to visualize data interactively, and to visualize the relationships between GWAS results from multiple cohorts or studies.

**Availability:** The **Assocplots** package is open source and distributed under the MIT license via GitHub (https://github.com/khramts/assocplots) along with examples, documentation and installation instructions.

## 1 Introduction

Advances in genotyping, sequencing, and phenotyping techniques have resulted in large quantities of genome-wide association studies (GWAS) data. The results of GWAS are commonly summarized and displayed on a Manhattan plot and a quantile-quantile (QQ) plot and help identify single nucleotide polymorphisms (SNPs) that are significantly associated with a given phenotype. Over the years, several standalone programs such as WGAViewer (Ge *et al.*, 2008), web applications (summarized in Zeigler *et al.*, 2015) such as LocusZoom (Pruim *et al.*, 2010) and packages (mostly written in R) such as qqman (Turner 2014) have been developed for producing these types of plots. Although dynamic, interactive visualization has recently become more widely adopted in genomic tools such as R/qtlcharts (Broman 2015) and LDlink (Machiela et al., 2015), it has not yet become a routine part of GWAS data analysis. Interactive data visualization not only allows clearer representation of multidimensional data, but also encourages a viewer’s engagement from simple data browsing to providing a platform for answering specific scientific questions, in ways that static data cannot. Here we present a python package for viewing GWAS results not only using classic static Manhattan and QQ plots, but also through an interactive extension which allows a user to visualize data interactively, for example: zoom into SNP dense regions, quickly obtain underlying details (e.g. SNP rs number or gene name, base pair position, p-value) by selecting a peak of interest, and visualizing the relationships between GWAS results from multiple cohorts or studies. For example our tool allows exploration of GWAS results from: (1) multiple phenotypes in a single group of individuals, (2) a phenotype measured among distinct cohorts, (3) expression quantitative trait loci measured across different tissues or cohorts, and (4) various experimental conditions such as before and after drug treatment. Thus, our tool makes it possible to browse multiple charts in real-time to better understand the relationships among groups.

## 2 Implementation and Features

**Assocplots** is implemented as a package for the Python programming language. Its basic functionality includes plotting interactive data visualization for viewing in the browser as well as static publication quality plots using matplotlib. Interactive visualization is implemented via a Python interactive visualization library, bokeh (http://bokeh.pydata.org/), that targets modern web browsers; and data wrangling is implemented with Numpy and Pandas scientific computing python libraries. All of these tools are open source. The use of python for this package makes it easily accessible to bioinformaticians, as it is one of the commonly used programming languages in the field. The package is designed to be used both in Jupyter notebooks (http://jupyter.org/) and in command line. Visualizing GWAS data in a web-based document (notebook), ensures data analysis reproducibility and makes it conveniently sharable with collaborators via online repositories such as GitHub. The **Assocplots** package is open source and distributed via GitHub under the MIT license. Below we present the package’s features and give an example of the plots (Fig. 1) using data from the Genetic Investigation of ANthropometric Traits (GIANT) consortium (Randall JC, *et al.*, 2013).

**Fig. 1.**
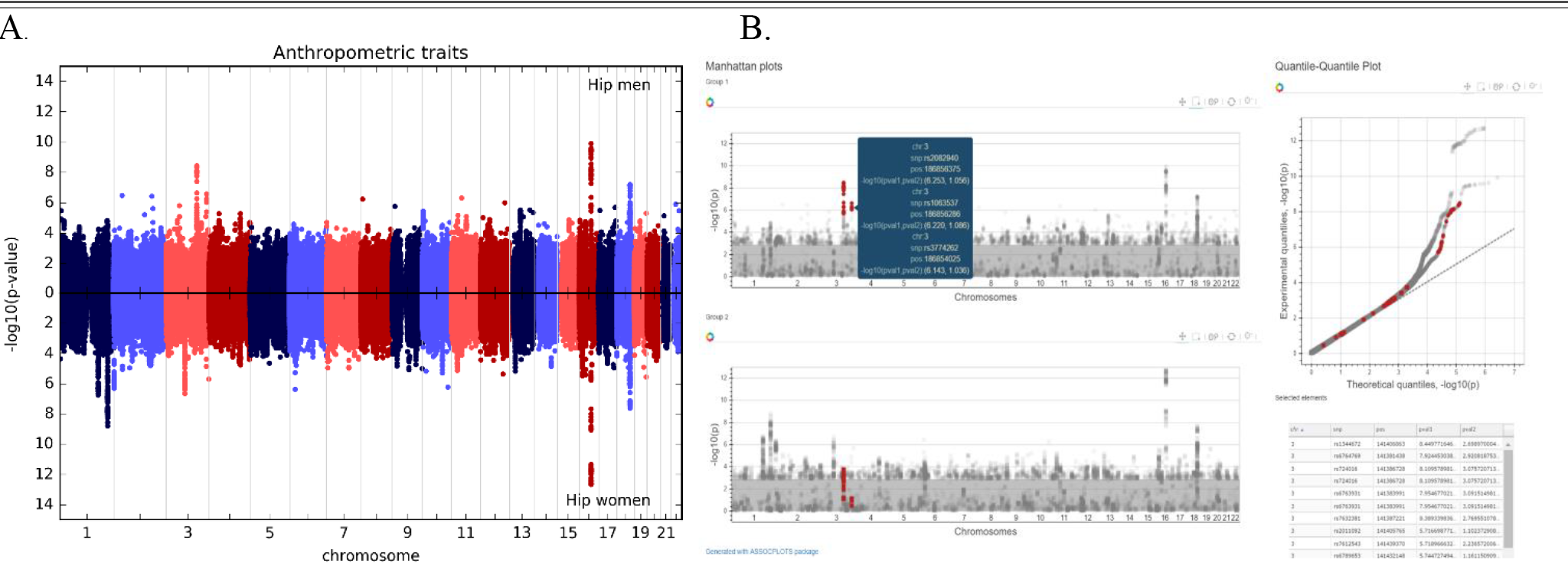
(A) Example of an inverted Manhattan plot generated using the static module and the data from the GIANT consortium, where each dot represents a single nucleotide polymorphism with the −log10(p-value) on the y-axis and chromosome and base pair position on the x-axis. (B) A static view of interactive Manhattan plots for two groups (same as in 1A). Selecting a group of SNPs in one of the groups highlights the same SNPs in the other group, and hovering over a dot shows the information associated with the SNP: chromosome, rs number, base pair position, and a −log10(p-value) for both groups. For the interactive version of this figure, see http://khramts.github.io/output.html.

### 2.1 Static module features

#### 2.1.1 Classic Manhattan plot

a. X-axis: chromosome and base pair (both numeric and alphabetical names, so various chromosome labeling (e.g. 1, 2, chr1, X) is acceptable)
b. Y-axis: Although −log10(p-value) is the most commonly used value for the y-axis, other values such as the effect size can be specified
c. Inverted Manhattan plot for two groups for easier visualization of peak differences **(Figure 1A).**

#### 2.1.2 Classic quantile-quantile plot

a. Multiple groups plotting: Multiple groups can be visualized on the same QQ plot for easier comparison.
b. Genomic Inflation Factor, λgc, calculation: In GWAS population substructure and cryptic relatedness among subjects can lead to spurious errors, and genomic control method is commonly used correct the underlying population stratification (Devlin et al. 1999). The static module can be used to calculate λgc.
c. Confidence Intervals (CI) estimation: The package allows to plot CIs for either the null distribution or the experimental data. When multiple groups are plotted, CI can be displayed for each group.

#### 2.1.3 Figure generation

Assocplots supports matplotlib plotting backends and thus can save figures in raster format (i.e. png and jpg) and vector format (i.e. pdf and ps).

### 2.2 Interactive module features

#### 2.2.1 Dynamic Manhattan and QQ plot

a. Info pop-up: Hovering over a point reveals information about the SNP/gene, such as the name (SNP rs number), chromosome, base pair location, and the statistic reported on the y-axis (−log10(p-value) or effect size) **(Figure 1B).**
b. Group comparison: Selecting a set of SNPs in one graph automatically highlights those same SNPs in the other graph (a different phenotype, population, condition, etc.) Additionally, a table is generated below the graphs, listing all the selected SNPs and information about those SNPs including the position, and the test statistic across groups.
c. Zoom-in and -out: Plotting many points on the same graph makes it difficult to discern one point from another, as it may be in a peak or in the lower portion of the Manhattan plot which often is densely packed. To overcome this issue, the plot can be zoomed-in using a mouse scroller when the mouse pointer is placed on the Manhattan plot.

#### 2.2.2 Visualization Sharing

Interactive plots can be saved as notebooks and self-contained html files that can be shared with colleagues via usual sharing platforms (GitHub, Dropbox, Google Drive, etc.) and opened in any modern web browsers on any operation system.

## 3 Limitations

In general, interactive visualization made through web browsers are limited by the number of objects they can smoothly display. In the current example **(Figure 1B)** we have selected the top (most significant) 1,000 SNPs in each of the two groups and are visualizing at most 2,000 dots if there is no overlap between those SNPs. To address this limitation, the package can be extended to a web application with dynamic data loading from a database/server. Dynamic data loading would allow a user to load SNP data in real time for a specific region of interest as the user zooms-in. By making this an open source package that is accessible via GitHub, we invite members of the scientific community to contribute and enhance the package’s capabilities.

## Acknowledgements

The authors thank Sasha Kaurov for advice on code implementation.

